# Metabolic and genomic characterization of two novel butyrate producing Lachnospiraceae isolated from swine feces

**DOI:** 10.1101/2023.12.14.571744

**Authors:** Hannah R. Watkins, Julian M. Trachsel, Shawn M. D. Bearson, Crystal L. Loving, Christopher L. Anderson

## Abstract

Commensal bacteria from the swine gut microbiome that can be isolated have numerous potential applications in the animal production industry, including mitigation of disease, improving performance, and promoting colonization resistance to human foodborne pathogens. Butyrate-producing bacteria are targets for next-generation probiotics and microbiome-engineering strategies because butyrate is a metabolite of central importance in large intestinal homeostasis and may augment colonization resistance to enteric pathogens. However, relatively few butyrate-producers from swine have been cultured and extensively characterized. Here, we describe the substrate utilization, metabolic profiles, and genomic features of two novel species that produce high concentration of butyrate *in vitro*, *Roseburia sp.* 831b and *Petralouisia sp.* 499, isolated from swine feces. The complete genomes illustrated versatility in carbon metabolism and unique carbohydrate-active enzymes not observed in other species of *Roseburia* and *Petralouisia* that encode a combination of glycosidic hydrolases and carbohydrate-binding modules involved in starch and pectin utilization. *Roseburia sp.* 831b fermented a broader range and more complex mono- and polysaccharides than *Petralouisia sp.* 499. Fecal and cecal metagenomes from eight-week-old pigs challenged with Salmonella revealed that *Roseburia sp.* 831b increased to detectable abundances in the swine hindgut in most animals at ∼63-70 days of age. Additionally, the abundance of *Roseburia sp.* 831b in fecal metagenomes correlated with fecal butyrate concentrations in the pigs fed a diet supplemented with a prebiotic resistant potato starch. Together, these findings highlight the probiotic potential and ecological niche in the swine gastrointestinal tract for two novel butyrate-producers.

**Importance:** Antibiotics have been important for swine production and management of enteric pathogens; however, the Veterinary Feed Directive limits the use of medically important in-feed antibiotics for production purposes. As a result, there is a need for alternatives to antibiotics. Butyrate-producing bacteria can improve colonization resistance to human pathogens within the swine gastrointestinal tract by reinforcing the intestinal barrier, increasing mucus production, and reducing local oxygen and pH levels. Here, we demonstrate the versatile substrate utilization and metabolic potential of two novel species isolated from swine that produce high butyrate concentrations *in vitro*. These findings will help develop strategies that increase the abundance of these species and other butyrate producers in the swine gut. Further, isolating and characterizing swine butyrate producers is necessary for controlled studies that provide a mechanistic understanding of how this functional group of bacteria promotes swine gut health and colonization resistance to bacteria of public health concern.

## Introduction

Swine are reservoirs for common human foodborne pathogens such as *Salmonella* (1), *Campylobacter* (2), and *Escherichia coli* (3, 4). These pathogens can colonize swine at various points of the production process, including through contaminated feed and water, as well as via contact with infected pigs, caretakers, and other vectors (5-7). Diverse measures have been implemented to control foodborne pathogens in swine production, such as strengthening biosecurity measures (8), establishing vaccination programs (9), improving sanitation (10), managing feed quality (5), practicing antibiotic stewardship (11), and promoting colonization resistance (12). Colonization resistance in the gastrointestinal tract can be mediated through a combination of direct (nutrient competition, antagonism through antimicrobial peptides, and inhibitory metabolites) and indirect (mucosal barrier, oxygen limitation, and modulation of the immune response) mechanisms (13, 14).

Butyrate is a short-chain fatty acid (SCFA) produced by intestinal commensal microbes. It can play a role in promoting colonization resistance against human enteric pathogens that are harbored in the swine intestinal tract (15-17). Butyrate can reduce the pH in the intestine, creating an acidic environment that is unfavorable for the growth of many pathogens (18). Additionally, butyrate is the preferred energy source for colonocytes (19-21), helping to strengthen the barrier function and prevent translocation of microbes across the gut lining (22). The oxidation of butyrate leads to a decrease in local oxygen levels, which in turn serves as a limiting factor for the growth of facultative anaerobic pathogens (23). Furthermore, butyrate promotes the production of mucus (24, 25), which acts as a physical barrier against pathogens. Butyrate also modulates the immune response, enhancing the activity of immune cells and promoting the production of antimicrobial peptides that can directly target and eliminate pathogens (19). By fostering a balanced microbial community, enhancing gut barrier function, and influencing the immune response, butyrate can play a role in bolstering the swine’s colonization resistance against pathogens in their intestinal tract and contributing to overall gut health.

Among important butyrogenic taxa of the gut microbiota, Lachnospiraceae is a family of phylogenetically and morphologically diverse obligate anaerobes within the *Clostridium* cluster XIVa (26). Many of these described species are exceedingly specialized in their ability to ferment a wide array of complex polysaccharides to SCFAs (27). Butyrate is the primary SCFA metabolite originating from Lachnospiraceae mediated digestion of recalcitrant substrates not directly utilized by the host (28, 29), contributing to maintained intestinal homeostasis, resistance to enteric pathogens, and enhanced epithelial barrier function (15, 30, 31). Lachnospiraceae populations are established in early life of the piglet, colonizing the intestinal lumen and increasing in abundance with age (32). Here, we characterize the substrate utilization, encoded metabolic potential, and ecological niche of two novel anaerobic butyrate-producing strains of Lachnospiraceae – *Roseburia sp.* 831b and *Petralouisia sp.* 499 – previously isolated from swine feces (33). Our characterizations provide insight into the probiotic potential of these two novel species.

## Results and Discussion

### Optimal culture conditions of Lachnospiraceae isolates aligned with the pig hindgut environment

Isolates 499 and 831b were previously cultured from the feces of a post-weaning piglet and healthy sow, respectively (33). Both isolates grew optimally at temperatures ranging from 37-39°C, in alignment with pig body temperature. Growth slowed noticeably at 35°C for both isolates (data not shown). At 32°C 831b failed to grow, while 499 grew with a delay of approximately three days to reach an OD^600^ value of 0.2. pH values ranging from 6.0-10.0 and 6.0-11.0 supported 499 and 831b growth, respectively, but the optimal media pH for cultivating both isolates was between 7.0-8.0 (data not shown).

Other Lachnospiraceae isolates range from short or elongated rods to cocci (34) and are often flagellated. While the flagellate status of other *Petralouisia* is undocumented, *Roseburia* has multiple subterminal flagella and species are predominantly motile (35, 36). Isolates 499 and 831b were observed with phase-contrast microscopy to be highly motile when cultured in RTY medium. Isolate 499 had flagellar tufts occurring in two sets, one subpolar and the other mid-cell. In contrast, 831b displayed subterminal flagella, sometimes appearing in multiple subpolar sets but consistently only at one end of the cell. Isolate 499 and 831b cells were rods, with 499 being straight to slightly curved and 831b straight and elongated by comparison (Figure S1); the cells of 831b further aggregated into chains in liquid culture after 24 h of incubation (data not shown). Cells of 499 measured 0.818-1.21 µm long and 0.385 µm in diameter, while 831b cells were 0.955-2.27 µm long and 0.442 µm in diameter (Figure S2). The most abundant fatty acids of 499 were C17:0 (38.7%), C14:0 (25.3%) and C15:0 (11.1%) and the major fatty acids for 831b were C17:0 (66.8%), C14:0 (7.3%), and C18:0 (6.2%) (Table S1). Collectively, 499 and 831b were motile anaerobes and the conditions best suited for culturing the isolates mirrored the environment of the adult pig hindgut (37, 38).

### Swine isolates 499 and 831b are novel species of *Petralouisia* and *Roseburia*

Previous 16S rRNA gene-based comparisons classified isolate 831b as *Roseburia* and 499 as a potentially novel genus or species, with *Roseburia hominis* as the closest related species (33). Later analysis suggested isolate 499 may belong within the genus *Pseudobutyrivibrio*, though bootstrap support for this assignment was low (39). We supplemented pre-existing short-read genomes with Oxford Nanopore sequencing to resolve the taxonomy and metabolic potential of isolates 831b and 499. Adding long-read sequencing data resulted in closed circular genomes for 499 (3.38 Mbp) and 831b (3.13 Mbp), with evidence of a large 243 kbp circular plasmid in 831b.

The resistance profile of *Petralouisia sp.* 499 consisted of chromosomally encoded *tetO*, *lnuC*, and *vanI* genes. The plasmid detected in *Roseburia sp.* 831b conveyed putative resistance to tetracycline and lincosamide (*lnuC*, *tetO*, and *tet*(40) genes). This plasmid was predicted to be non-mobilizable. Additionally, bacteriocin resistance genes were identified in both isolates, which may contribute to the ability of strains 499 and 831b to effectively colonize the gut environment (40).

Isolate 499 was placed within the newly described genus *Petralouisia* (type strain *Petralouisia* muris) (Figure 1) (41), currently designated as “g *14-2”* in the Genome Taxonomy Database (GTDB) (42). *Petralouisia* resides in the family *Lachnospiraceae,* phylogenetically between *Eubacterium* and *Pseudobutyrivibrio* (41). Isolate 831b was confirmed through phylogenetic analysis to belong to the genus *Roseburia* (Figure 1). Both isolates 499 and 831b represented novel species, as they shared low average nucleotide identity (ANI) (<95%, the proposed definition to delineate species through genomic sequences (43)) and alignment fraction to other members of their respective genera. The genomes that shared the highest ANI with 499 were *Petralouisia sp00403255* (isolated from mouse feces, 85% ANI) and *Petralouisia sp012519695* (isolated from crab-eating macaque feces, 85% ANI, Figure 2A). The GC content of 499 was 38.2% mol as compared to 44% for the type strain *Petralouisia muris.* To date, isolate 499 is the only species of *Petralouisia* cultured from the swine gastrointestinal tract. Isolate 831b shared 86.7% sequence identity with *R. hominis*, followed by 86.2% sequence identity with *R. intestinalis* (Figure 2B). The GC content of 831b was 40.7%, in alignment with other reference *Roseburia* strains, which ranged from 41.4-47.4% (36). A phylogenetic tree based on 16S rRNA gene sequences of *Roseburia* and *Petralouisia* isolates similarly distinguished isolates 831b and 499 from other major clades of their respective genera (Figure S3). Given the novelty and butyrate producing capacity of *Petralouisia sp.* 499 and *Roseburia sp.* 831b isolates, we further investigated the genomic and biochemical properties of the isolates.

**Figure 1.**
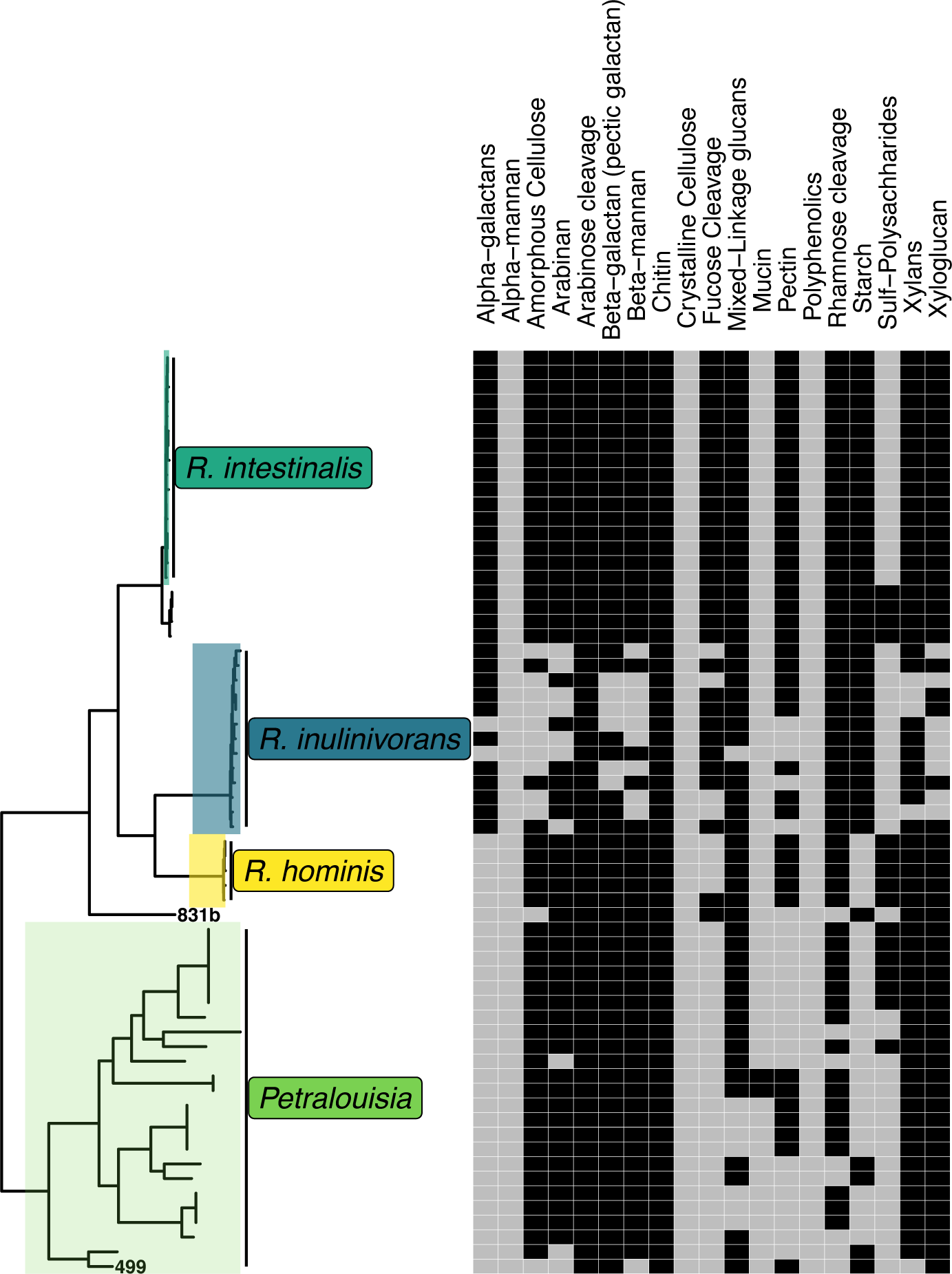
Substrate-resolved assessment of carbon metabolism for *Roseburia* and *Petralouisia* isolates, including 499 and 831b (dark squares indicated predicted activity and light gray squares denote absence). CAZymes were linked to broader substrate categories with DRAM to predict substrate utilization. The phylogenetic tree was constructed based on the alignment of single copy core bacterial genes and is rooted at the mid-point.

**Figure 2.**
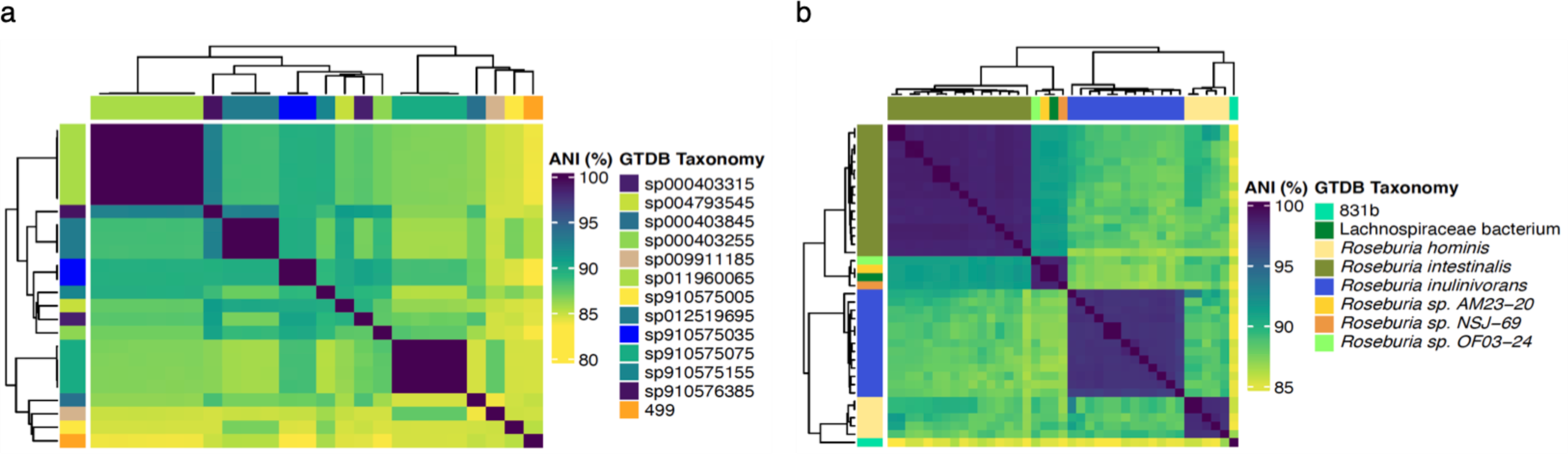
Heatmap and dendrogram depicting the pairwise ANI values between *Petralouisia* (a) and *Roseburia* (b) isolate genomes. Strains with ANI values >95% (suggested as a threshold to delineate bacterial species based on genomic content (43)) are represented in darkest shade of blue

### Novel Lachnospiraceae isolates produced high concentrations of butyrate *in vitro*

Synthesis of SCFAs occurs as a result of bacterial fermentation of polysaccharides, lignin, oligosaccharides, and resistant starches that are inaccessible to host enzyme hydrolysis and reach the hindgut intact (28). Butyrate in particular fuels large intestinal epithelial cell metabolism and integrity through the oxidation of the compound (44) and may also suppress pathogen colonization through various proposed mechanisms (45, 46). Four major microbial pathways lead to the production of butyrate: 4-aminobutyrate/succinate, glutarate, lysine, and acetyl-CoA. All these pathways converge at the point of butyryl-CoA generation, thereafter differing in the final enzymes and co-substrates that are used to yield butyrate. The acetyl-CoA pathway of butyrate production is most common among gut microbes, the terminal step of which can be catalyzed by either butyryl-CoA:acetate CoA transferase (*but* gene) or butyrate kinase (*buk* gene). Alternative transferases can be encoded in rare cases (47). These genes, regarded as biomarkers for butyrate production, have been screened for in many studies, although in pigs and other hindgut fermenters, *but* is most abundant (48, 49). Amino acids (glutarate, lysine, 4-aminobutyrate) or inputs from cross-feeding between other gut bacteria, particularly species that produce acetate, which are required by *but* (47), can fuel butyrate production.

It was previously confirmed through PCR that 499 and 831b possess *but*, and activity was verified *in vitro* through cloning and transforming the genes into *Escherichia coli* (49). We annotated and extracted *but* genes from reference *Roseburia* and *Petralouisia* isolate genomes to investigate the divergence in sequences amongst the two genera (Figure 3). This analysis revealed a clade of *Petralouisia* with *but* genes more closely related to *Roseburia*, similar to the evolutionary relationships observed amongst the 16S rRNA genes of these strains (Figure S3). Further, several strains of *Petralouisia* had multiple copies of *but* and putative *buk* genes. Genomes possessing *but* and *buk* have been documented previously (50), including in *Roseburia inulinivorans* LI-83, which was also identified in our analysis. Additional *in vitro* testing would be necessary to determine whether *buk* is expressed and functional in cases when both terminal enzymes are encoded in the genome. It is unclear why genomes would encode both *but* and *buk* genes; however, given that *but* is involved in acetate consumption and *buk* is not, we speculate that *buk* could be preferentially used as the terminal enzyme when acetate is not as readily available.

**Figure 3.**
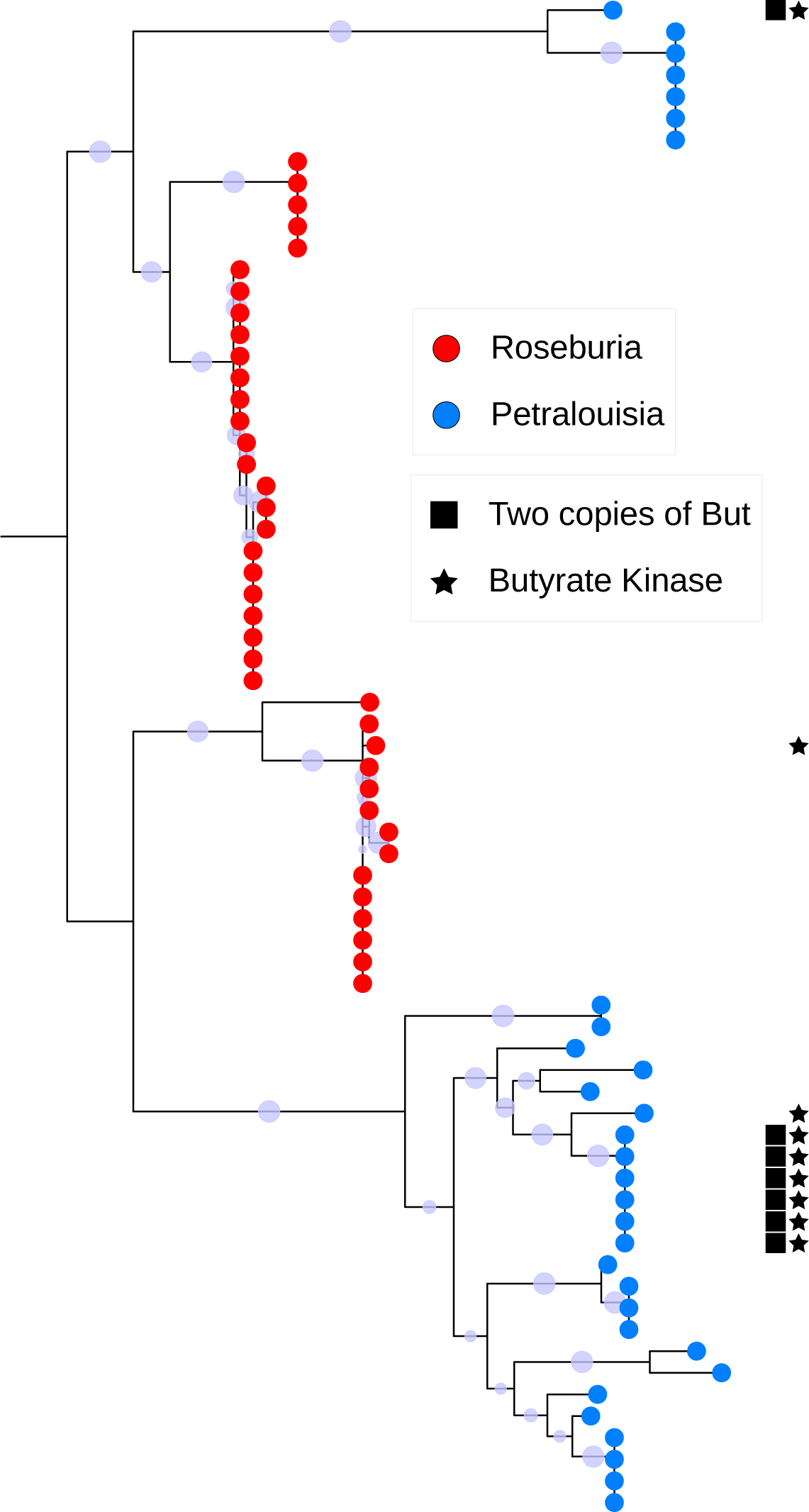
Maximum likelihood tree based on the alignment of butyryl-CoA:acetate CoA transferase (*but*) amino acid sequences from *Roseburia* and *Petralouisia* isolate genomes. Additionally, the genomes that encoded multiple copies of *but* or putative butyrate kinase (*buk*) genes are denoted. Bootstrap values are shown on internal nodes and the tree is rooted at the midpoint.

The SCFA fermentation products of 499 and 831b were quantified via high-performance liquid chromatography (HPLC). Isolate 499 produced 12.2 mM butyrate, whereas isolate 831b yielded 22.4 mM butyrate from the fermentation of glucose (Table 1). SCFA profiles have only been reported for *Petralouisia muris*, the type strain of the genus, which produced approximately 8 mM of butyrate through the utilization of acetate when cultivated in Yeast Casitone with Fatty Acids (YCFA) medium (41). The SCFA profile of 831b was consistent with reports of other *Roseburia* species, wherein butyrate is the major end-product of fermentation (36, 51-53) (Table 1). Other SCFAs produced by members of *Roseburia* are formate and lactate; however, our SCFA panel did not include lactate. SCFA production profiles also depend on the substrate used for fermentation – that is, fermentation of one substrate may lead to more butyrate production than another. Therefore, though not inherently comparable, only a single strain of *Roseburia faecis* demonstrated a greater concentration of butyrate production (24 mM) on M2GSC medium (glucose, cellobiose, soluble starch as carbon sources) than isolate 831b (22.4 mM with glucose) (36).

**Table 1.** Key phenotypic characteristics of isolates 499, 831b, and selected reference species of *Petralouisia* and *Roseburia*. Data for reference species was obtained from Afrizal *et al.* (*P. muris*) (41), Wylensek *et al.* (*R. porci*) (53), Duncan *et al.* (*R. faecis, R. hominis, R. inulinivorans*) (36) Duncan *et al.* (*R. intestinalis*) (51), and Stanton and Savage (*R. cecicola*) (52). NR, Not reported.

| Characteristic | 499 | 831b | *P. muris* | *R. porci* | *R. faecis* | *R. hominis* | *R. inulinivorans* | *R. intestinalis* | *R. cecicola* |
| --- | --- | --- | --- | --- | --- | --- | --- | --- | --- |
| Isolation source | Pig feces, sow | Pig feces, post-weaning piglet | Mouse cecal/colon contents | Pig feces | Human feces | Human feces | Human feces | Human feces | Mouse cecal mucosa |
| Cell shape | Rod, slightly curved | Rod | NR | Rod, slightly curved | Rod, slightly curved | Rod, slightly curved | Rod, slightly curved | Rod, slightly curved | Rod, slightly curved |
| Cell size (µm) | 0.4x0.8-1 | 0.4x1-2 | NR | 1 (length) |  | 0.5x1.5-5 | 0.5x1.5-5 | 0.4-0.5x1-5 | 0.5x2.5-5 |
| Motility | + | + | NR | + | + | + | + | + | + |
| G+C content (mol%) | 38.2 | 40.7 | 44.0 | 41.7 | 42.0 | 47.4 | 41.4 | 42.6 | 42.3 |
| Substrate utilization: |  |  |  |  |  |  |  |  |  |
| D-glucose | + | + | NR | + | + | + | + | + | + |
| D-lactose | + | + | NR | NR | NR | NR | NR | NR |  |
| D-sucrose | + | + | NR | NR | - | - | + | + | + |
| D-maltose | + | + | NR | NR | + | + | + | + | + |
| Salicin | - | - | NR | NR | NR | NR | NR | NR | NR |
| D-xylose | - | + | NR | NR | + | + | - | + | + |
| L-arabinose | - | - | NR | NR | - | + | - | + | NR |
| D-cellobiose | - | + | NR | NR | + | + | + | + | + |
| D-mannose | + | - | NR | NR | NR | NR | NR | NR | NR |
| D-raffinose | - | + | NR | NR | + | - | - | + | + |
| Fructose | - | + | NR | NR | + | + | + | + | NR |
| Esculin hydrolysis | - | + | NR | NR | NR | NR | NR | + | - |
| Gelatin hydrolysis | + | - | NR | NR | NR | NR | NR | NR | NR |
| Indole formation | - | - | NR | NR | NR | NR | NR | NR | NR |
| Urease | - | - | NR | NR | NR | NR | NR | + | NR |
| Catalase | - | - | NR | NR | - | - | - | + | - |
| Fermentation products (mM): |  |  |  |  |  |  |  |  |  |
| Formate | - | - | - | + (5.1) | + (9.2-15.7) | + (6.9-7.6) | + (4.4-4.7) | + (6.9-8.4) | - |
| Propionate | - | - | - | - | - | - | - | - | - |
| Lactate | NR | NR | - | + (8.8) | + (0-2.3) | + (2.2-0.1) | + (2.1-7.1) | + (0-7.1) | - |
| Butyrate | + (12.2) | + (22.4) | + (8) | + (12.3) | + (13.9-24) | + (16.8-20) | + (6.5-7.3) | + (16-21.3) | + (NR) |
| Succinate | - | - | - | - | - | - | - | - | - |

### Carbohydrate-active enzyme profiles revealed differences in substrate specificity across *Roseburia* and *Petralouisia*

Various polysaccharides are utilized and fermented by butyrate producers. Butyrate production can be stimulated directly by dietary components or through metabolic cross-feeding. As a result, diet can play a prominent role in dictating the diversity of butyrate-producing bacteria in the large intestine. Given the range of polysaccharides that butyrate-producing bacteria can utilize and the importance of carbohydrate metabolism in establishing the ecological niche of these organisms in the intestinal environment, we next investigated the carbohydrate metabolic versatility of 831b and 499 in the context of other members of *Roseburia* and *Petralouisia*.

Carbohydrate-active enzymes (CAZymes) facilitate the degradation of oligosaccharides and polysaccharides. CAZymes can be classified into families of glycoside hydrolases (GH), carbohydrate esterases (CE), glycosyl transferases (GT), polysaccharide lyases (PL), redox auxiliary activities (AA), and carbohydrate-binding modules (CBM) (54). Our analysis of carbohydrate metabolic potential (Figure 4) suggested all *Roseburia* and *Petralouisia* species lack genomic evidence for crystalline cellulose, alpha-mannan, and polyphenolic cleavage. Conversely, all genomes included in our comparisons possessed various GHs and CEs contributing to arabinose and chitin cleavage, as well as amylomaltase/4-alpha-glucanotransferase (GH77, EC 2.4.1.25), an enzyme with starch-modifying functions (55). GH94, a family attributed to the degradation of amorphous cellulose, was detected in all members of *Roseburia* and many *Petralouisia* species; however, *Roseburia* had a higher copy number of GH94 genes on average. *Roseburia* also appeared equipped for saccharification of alpha-galactan and fucose moieties, characteristics not annotated in species of *Petralouisia*. We identified carbohydrate-binding modules (CBM48) with glycogen-binding functions in both genera’s genomes. Isolates 499 and 831b encoded CBM26 and CBM41, families associated with starch-binding (56). Only *Petralouisia sp012519695* also harbor these CBMs. CBM41 is a protein domain with residues found predominantly in bacterial pullulanases (57).

**Figure 4.**
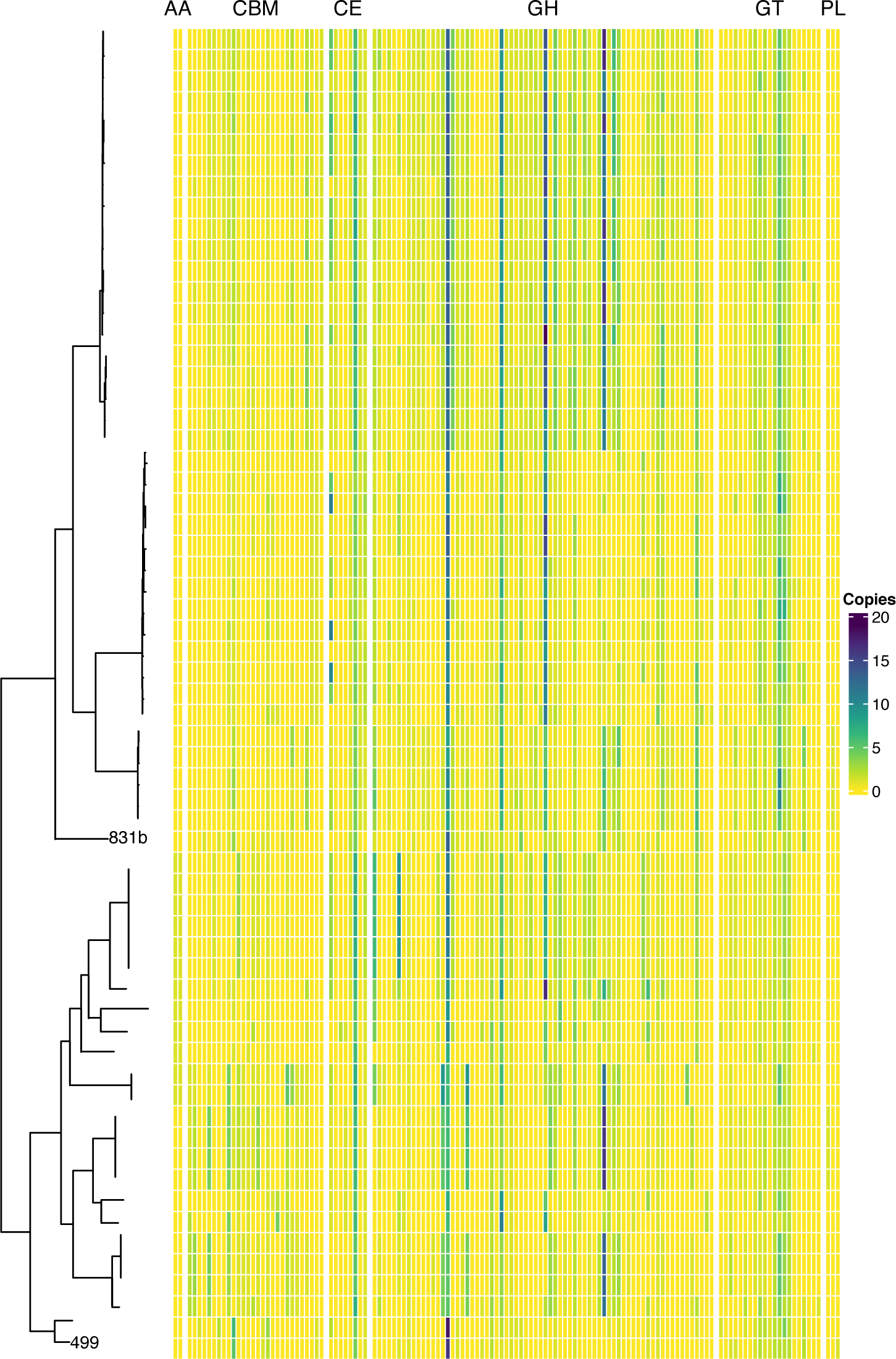
Heatmap of the carbohydrate active enzyme gene content for *Petralouisia* and *Roseburia* isolates grouped by CAZyme families. The CAZyme families refer to auxiliary activities (AA), glycoside hydrolases (GH), glycosyl transferases (GT), polysaccharide lyases (PL), carbohydrate esterases (CE), and carbohydrate binding modules (CBM). The color gradient reflects gene copy number in each genome. The phylogenetic tree was constructed based on the alignment of single copy core bacterial genes and is rooted at the mid-point.

Annotation of CAZyme gene content revealed a unique pattern of carbon metabolism in isolate 499, as genes encoding pyruvate formate-lyase activating enzyme, citrate (Re)-synthase, various GHs, GTs, and CBMs for starch molecules being among the list of CAZymes shared with select members of *Petralouisia*. The CBM35 family of approximately 130 residues with varying substrate specificities (uronic acids, neutral sugars) (58) was found only in the genomes of *Petralouisia* species compared to *Roseburia* but was absent in isolate 499 and four other genomes. Certain GHs and CEs indicative of pectin oligo cleavage were widespread across *Roseburia* (though not in 831b) and not in *Petralouisia*. However, of all the *Petralouisia* and *Roseburia* genomes included in our analysis, 499 alone harbored pectate lyase (PL9, EC 4.2.2.2), an enzyme responsible for the degradation of pectic plant cell wall constituents via hydrolysis of α-1,4-glycosidic bonds. Pectate lyases differ from pectin lyases in both active site specificity to substrate and pH at which enzymatic function is optimal (59-61). Pectate lyases favor elevated pH values by comparison, which is perhaps indicative of the ecological niches isolate 499 may occupy in the pig gastrointestinal tract. Provided PL9 is important to the maintenance of 499 *in vivo*, peak enzymatic function could be achieved following the pH gradient that exists from small intestine to terminal the ileum before dropping significantly in the cecum and gradually increasing from colon to rectum, where highest pH values would likely be observed (62). According to this reasoning, highest abundances of isolate 499 would likely be observed in the cecum or colon compared to small intestinal compartments. Future work could focus on confirming the growth and optimal conditions for the growth of 499 on pectin substrates relative to closely related strains.

Most *Roseburia* genomes harbored a shared array of CAZyme genes, though several traits were distinctive. For instance, all *Roseburia* species except for 831b possessed the GH29 family involved in fucose oligo cleavage. Among the class of CBMs, only the 831b genome contained CBM20 and CBM56, described as having granular starch and β-1,3-glucan binding functions, respectively (63, 64). Isolate 831b, *Petralouisia sp000403255*, and *Petralouisia sp000403315* could also be distinguished by the absence of sucrose synthase (GT4, EC 2.4.1.13). The presence of genes encoding β-mannanase (GH113), α-L-fucosidase (GH95), and β-1,3-glucan phosphorylase (GH161) was not unique to 831b, but also not widely distributed across *Roseburia*. Many *Roseburia* genomes, including 831b, contained the gene encoding amylo-α-1,6,-glucosidase (GH133, EC 3.2.1.33), an enzyme involved in starch cleavage. Interestingly, 499 also possessed this genomic feature, but it was not observed in other *Petralouisia* genomes in our analysis. Following a similar pattern, CBM34 was detected in select members of *Roseburia* (not 831b or other 14-2 isolates) and isolate 499. This CAZyme has a putative starch-binding function as well (65). Together, these findings highlight the potential role of 831b and 499 in starch binding and hydrolysis in the swine hindgut. Metabolism of starch may convey a competitive advantage to these isolates in specific dietary contexts.

Genes associated with starch utilization are common across Lachnospiraceae (66), though there is a large degree of variability between genera and species. Dietary resistant starch (RS) interventions to manipulate gut microbial communities have induced an increase in butyrate producing Lachnospiraceae, including *Roseburia* (67, 68). Taxa able to utilize starch are highly specialized in this function, and different RS classes are associated with varying trends in gut microbiota composition (69). Feeding RS type 2 to humans, pigs, and rodents increases the abundance of *Roseburia* (67, 69, 70). RS2 is resistant to enzymatic cleavage due to its ungelatinized, granular structure. Sources of RS2 include raw potatoes, green bananas, and high-amylose corn starches (71, 72). While *Roseburia* may respond to such feed amendments, a description of their functional role in the complex network of starch degradation is needed.

Of note, neither 499 nor 831b isolates possessed GHs associated with mucin degradation, and only two other *Petralouisia* species were predicted to degrade these specific glycan linkages. Not encoding genes for mucin degradation is potentially favorable for probiotic species, as in certain contexts mucin degradation may damage the host epithelium (73, 74). The absence of CAZymes specific to host sugars suggests 499 and 831b may not colonize the mucosa of the gastrointestinal tract. Despite the lack of genomic evidence to support mucin degradation, exploring *in vitro* ability of host sugars to be the sole carbon source to support growth would be beneficial. The proportion of mucin-degraders among butyrate-producing species is unclear; however, in a study that isolated fifteen unique butyrogenic species from the pig gut, 20% grew in a medium containing mucus as the sole carbon source (33).

### Substrate and biochemical assays indicate isolates 499 and 831b utilize different sugars

Given the genomic predictions above, we aimed to assess the utilization of a subset of common monosaccharides, disaccharides, and polysaccharides *in vitro* (Table 1). Isolate 831b exhibited growth improvement for all sugars tested except D-mannose, with robust growth (OD^600^ ≥ 0.8) on cellobiose, maltose, sucrose, and D-raffinose; moderate growth (OD^600^ ≥ 0.4) on D-fructose, and weak growth (OD^600^ < 0.4) on L-arabinose and D-glucose. Meanwhile, isolate 499 demonstrated significant growth improvement on D-glucose, and weak growth on D-sucrose, D-lactose, D-maltose, and D-mannose. In addition, the API diagnostic kits demonstrated isolate 499 can hydrolyze gelatin, and produced acid from glucose, mannose, lactose, sucrose, and maltose. In contrast, isolate 831b hydrolyzed esculin, and produced acid from a broader range of sugars, including xylose, cellobiose, and raffinose. Neither of the isolates produced indole, and both were catalase and urease negative. These assays suggest 831b may be able to degrade more complex sugars (disaccharides and the trisaccharide raffinose) compared to 499, with the diminished ability to metabolize some monosaccharides. Paired with the genomic predictions of substrate utilization, the differences in carbohydrate metabolism between 831b and 499, and more broadly between *Roseburia*, and *Petralouisia*, may dictate their ecological niches in the swine gastrointestinal tract depending on dietary compounds available in the hindgut.

### Spore formation for isolates 499 and 831b remains inconclusive

Many gut bacteria utilize sporulation to enhance survival outside the host and promote transmission (75). It is also advantageous for anaerobic probiotic species to form spores due to the stability, ease of delivery, and ability to traverse the gastrointestinal tract. Lachnospiraceae is variable in its spore-forming content, with *Sporobacterium* being the most notable example (34, 76). Further, a recent analysis suggested the majority of butyrate producing bacteria in the human gut may be potential spore formers (77). Traitar (78), a tool for predicting microbial phenotypes from genomic sequences, suggested 499 may be capable of spore formation but not 831b. Common strategies to identify spore-formers employ chemical and environmental manipulations to isolate resistant spores by eliminating vegetative cells while retaining spores prior to direct culturing or addition of a germinant, such as bile acids (79). Ethanol shock followed by bile acid and oxygen exposure techniques did not yield colony-forming units for either strain after incubation at 39°C for 72 h. Yet, the two isolates could be former spore formers (an estimated 18% of Lachnospiraceae are former spore formers (75)) or the conditions employed may have been insufficient to induce sporulation *in vitro*. Given the extremely oxygen-sensitive nature of many butyrate-producing gut microbes, it would be valuable to experimentally validate the sporulation status of this diverse functional group of bacteria, and a potential key for considering an organism as a probiotic.

### The abundance of 831b correlated with fecal butyrate concentrations in pigs fed a diet amended with resistant potato starch

To investigate the potential ecological niches of these strains, we sequenced swine fecal and cecal metagenomes from a previously published study investigating the impact of a commercially available resistant potato starch (RPS) product on *Salmonella* shedding in pigs following experimental inoculation (80). In brief, pigs were fed either a control or RPS amended diet (5%) for 28 days following weaning, and subsequently challenged with *Salmonella* I 4,[5],12:i:-. Samples were collected to assess shedding dynamics for 21 days post-inoculation (DPI). In that study, pigs fed RPS had significantly lower cumulative fecal shedding of *Salmonella*. The decrease in *Salmonella* shedding could rely on an increased abundance of microbes capable of RPS degradation and bacterial cross-feeding interactions that would lead to the synthesis of specific SCFAs, including butyrate.

We conducted metagenomic sequencing on fecal samples from pigs at 0, 2, 7, 14, and 21 DPI as well as on cecal samples collected at necropsy at 21 DPI. Mapping of the metagenomic reads to the strain’s complete genomes and implementing a detection criterion of 0.5 genomic coverage breadth to assess the presence of a strain in a metagenome (81, 82) demonstrated 831b could be reliably detected in more fecal samples from both Control and RPS diets at 21 DPI (∼70 days of age) than at any other time point (Figure 5). Applying the same thresholds, 831b could not be detected in the cecal metagenomes. Together, this suggested the 831b strain may colonize or increase in abundance in the hindgut in later stages of the nursery phase regardless of diet. Isolate 831b was cultured from the feces of a sow, also suggesting 831b is detected in later life. There was no significant difference in the covered fraction of the 831b genomes between the Control diet and animals fed a diet with RPS at 14 and 21 DPI (∼63 and ∼70 days of age, respectively; Figure 5; Wilcoxon Rank Sum Test, *P* > 0.05).

**Figure 5.**
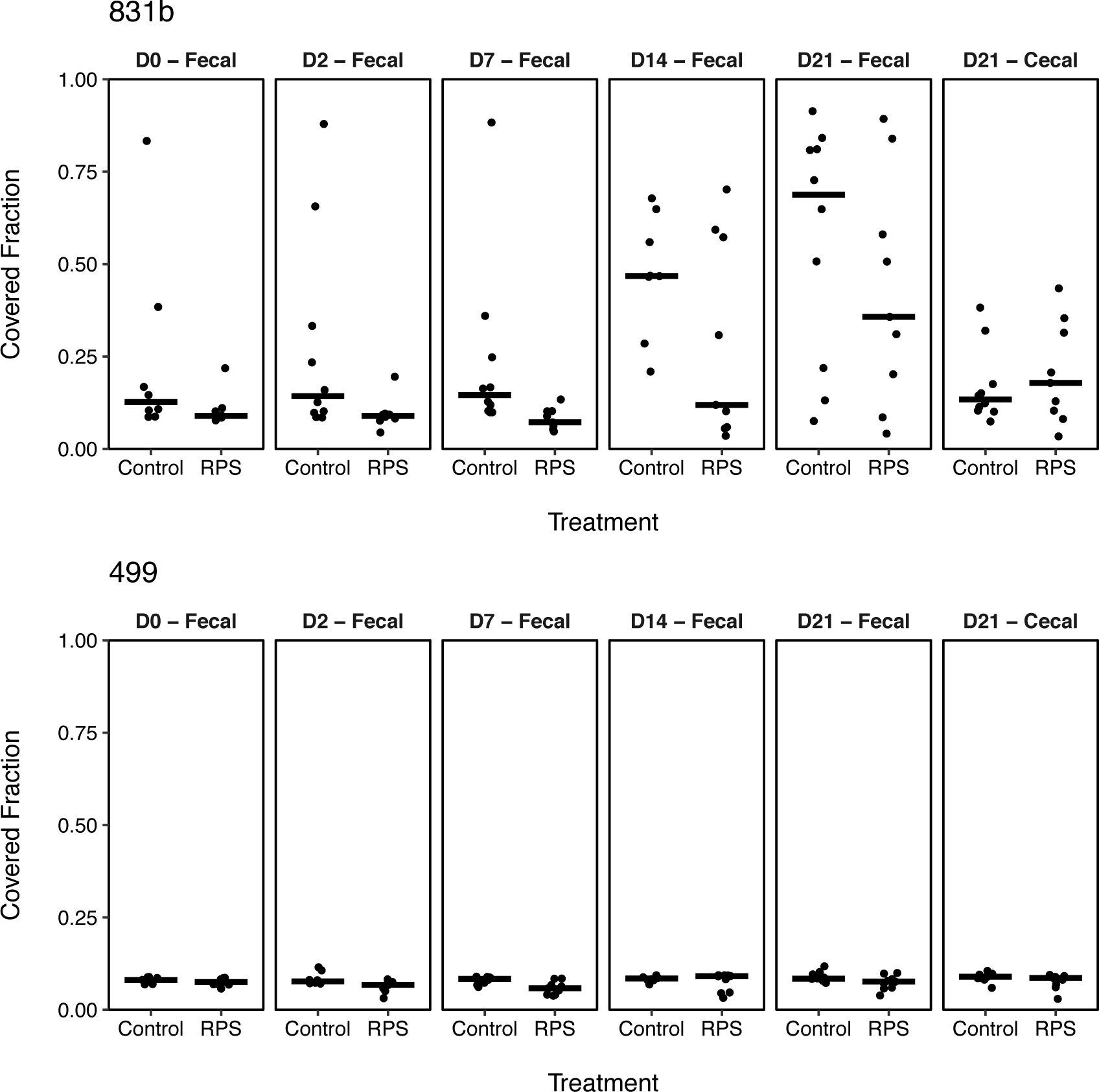
Fraction of the 831b (a) and 499 (b) genomes with at least 1X coverage in metagenomes from samples collected at different time points after *Salmonella* challenge in pigs feed a control diet or a diet supplemented with RPS. Bars indicate the median covered fraction. A coverage breadth of 0.5 would indicate at least 50% of the genome had at least 1X coverage and has been suggested as a threshold for the assessing the presence of a strain in a metagenomic dataset through read mapping (60, 61).

In the *Salmonella* challenge study, cecal butyrate concentrations tended to inversely correlate with cecal *Salmonella* shedding, though this trend was not observed in the fecal samples (80). However, others have associated decreased *Salmonella* shedding with butyrate concentration in the hindgut and also with the feeding of sodium butyrate (17). As a result, given 831b was a potent butyrate producer *in vitro* (Table 1), we investigated the relationship between the genomic coverage and abundance of 831b with fecal butyrate concentrations measured at 14 and 21 DPI, days in which 831b was reliably detected (Figure 5). There was a significant linear relationship between the coverage breadth (Figure 6a; linear regression, R^2^ = 0.32, *P* = 0.015) and reads per kilobase per million mapped reads (RPKM; Figure 6b; linear regression, R^2^ = 0.34, *P* = 0.012) of 831b and fecal butyrate concentrations in pigs fed the RPS diet. Supporting this finding, fecal butyrate concentrations were significantly higher for samples in which 831b had a coverage breadth greater than 0.5 (Figure S4; Wilcoxon Rank Sum Test, *P* = 0.010). The relationship between fecal butyrate concentration and 831b coverage and abundance was not confounded by the sampling date. This association was not observed in samples collected from pigs fed the Control diet on these same days (Figure 6; Wilcoxon Rank Sum Test, *P* > 0.05). Additionally, there was no significant relationship between the coverage breadth or abundance of 831b and *Salmonella* fecal shedding on these days for pigs fed the Control or RPS amended diet (*P* > 0.05).

**Figure 6.**
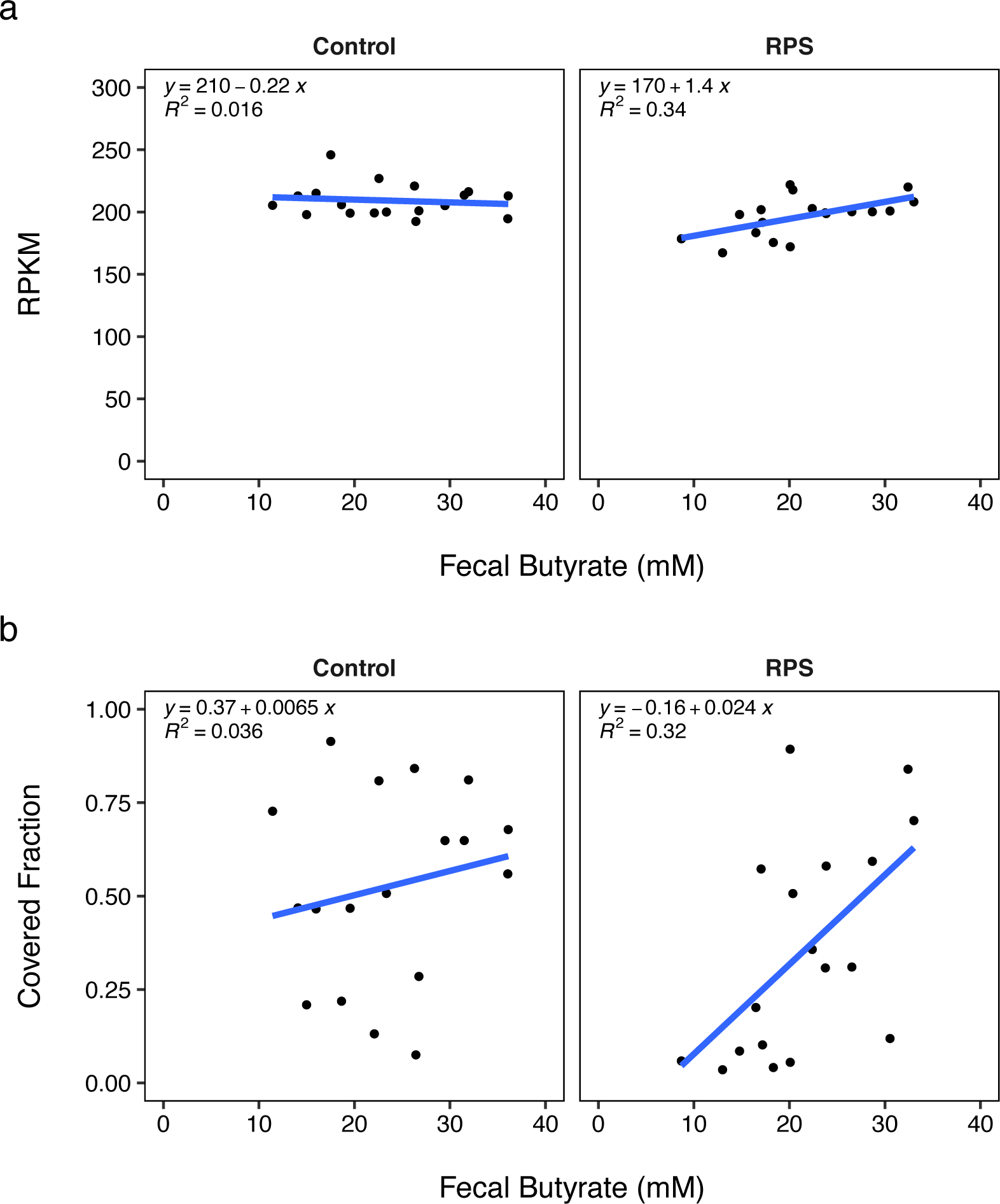
Relationship between fecal butyrate concentration and the RPKM (a) and coverage breadth (i.e., fraction of the genome with at least 1X coverage) of 831b in samples collected at 14 and 21 days after *Salmonella* inoculation in pigs feed a control diet or a diet supplemented with RPS. The RPKM and covered fraction values were calculated through the mapping of metagenomic reads from the corresponding fecal samples to the 831b isolate genome. A linear regression was fit to the data and the regression equation and correlation coefficient are depicted.

It is important to consider these associations were detected in animals challenged with a large dose of *Salmonella* in a controlled, experimental setting, which could alter the swine gut microbiota relative to pigs in a production environment. As a result, the colonization of 499 and 831b strains and relationships between 831b abundance and fecal butyrate concentrations should be extrapolated to animals in production settings with caution. The association between 831b abundance and fecal butyrate concentration was not likely due to the butyrate production of 831b alone, but rather that 831b may be part of a microbial consortia in the swine hindgut that helps promote butyrate production in certain ecological contexts, such as the environment created during the feeding of RPS. Others have demonstrated how co-culturing can be used to increase the abundance of butyrogenic bacteria and butyrate concentration (83, 84). Therefore, it is important to gain a better understanding of the microbial interactions that can increase butyrate production *in vivo*. We believe these findings highlight why some probiotics, and in general feed additives, have varied effect (85, 86) - the context of the varied microbial community members, diet, and broader intestinal environment likely plays a significant role in shaping the success of products designed to manipulate specific members of the gut microbiome. Future work may focus on unraveling the cross-feeding networks involved in the degradation and fermentation of RPS in the swine hindgut.

Strain 499 did not have a coverage breadth greater than 0.5 in any fecal or cecal metagenomes, suggesting this specific strain was not readily detectable in the samples from the *Salmonella* challenge study at the given sequencing depth (Figure 4). Reads that aligned to 499 may have been from other strains of the same species or aligned to conserved genomic regions. Methods with better sensitivity than metagenomic sequencing, such as quantitative PCR with primers targeting 499, could be used to detect and quantify the abundance of the strain in different compartments of the swine intestinal tract and explore potential relationships between 499 abundance and dietary components or SCFA production.

## Conclusions

The current study provides fundamental knowledge regarding the ecological niche, physiology, and genomic attributes of two new species of Lachnospiraceae that produce high concentrations of butyrate *in vitro*, *Roseburia sp.* 831b and *Petralouisia sp.* 499. We demonstrated the two isolates utilized a range of mono-and polysaccharide substrates and possessed carbohydrate-active enzymes not observed in other species of *Roseburia* and *Petralouisia*. General understanding of how interactions between different species and environmental nutrient availability influence bacterial butyrate production is limited. As a result, further work is needed to understand the regulation of butyrate metabolism in the pig intestinal tract. Isolating and characterizing *Roseburia sp.* 831b and *Petralouisia sp.* 499 is progress towards that goal. Future experiments could focus on methods to increase the abundance of 499 and 831b in the swine intestinal tract and evaluate the subsequent associations between colonization of 499 and 831b with shedding of foodborne *Salmonella* isolates. With further insights into 499 and 831b’s metabolism and co-culturing experiments, we hypothesize the abundance of these isolates could be increased through feed amendment strategies. Using specific polysaccharides as dietary additives, or through other microbial engineering strategies, such as designing microbial consortia in the context of the right prebiotic, butyrate production may be promoted in the swine intestinal tract.

## Methods

### Bacterial isolation and cultivation

Isolates 499 and 831b were initially cultured from pig feces in a targeted effort to identify butyrate-producing bacteria from the swine intestinal tract and were cultivated at that time in a depleted rumen fluid medium (DRF7) (33). Throughout this study, the isolates were maintained on a rumen fluid tryptone-yeast (RTY) medium (87) with resazurin as a redox indicator. All culture work was performed in a Coy Anaerobic Chamber (Coy Laboratory products) with atmospheric conditions of 80% N^2^, 10% CO^2^, and 10% H^2^ unless otherwise indicated. Glycerol stocks were prepared using equal volumes of 50% glycerol solution and overnight bacterial culture in 2 mL screw-cap cryovials for long-term storage at -80°C.

### Long-read sequencing and genomic analyses

To supplement existing Illumina MiSeq 16S rRNA gene sequencing data and obtain complete 499 and 831b genomes, DNA was isolated using PureLink’s genomic DNA extraction kit (Invitrogen) according to manufacturer instructions. The DNA libraries were prepared using the SQK-RBK004 Rapid Barcoding Kit (Oxford Nanopore), following the standard protocol. Individual Flongle Flow Cells (FLO-FLG001) were used in a MinION device and run for 24 hours. Quality checks of short- and long-read data were performed using fastp (v0.23.2) and Filtlong (v0.2.1) (88, 89). Illumina and Nanopore reads were assembled with Unicycler (v0.4.9) (88, 89), and classified with the Genome Taxonomy Database Toolkit (GTDB-Tk, v2.1.0) (42). Average nucleotide identity (ANI) values were calculated and visualized with pyani (v0.2.12) (90).

Bacterial ribosomal RNA predictor (barrnap, v0.9) was used to extract 16S rRNA gene sequences from the cultured isolates and reference *Roseburia* and *Petralouisia* genomes identified in GTDB (GTDB R207, metagenome-assembled genomes were not included) (91). The 16S rRNA gene sequences from all isolate genomes were aligned with ssu-align (v0.1.1) (92). IQ-TREE (v2.2.0) was used to construct a full-length 16S rRNA based maximum-likelihood tree which was then visualized with Interactive Tree of Life (iTOL, v5) (93-95). Functional annotations were assigned with the Distilled and Refined Annotation of Metabolism (DRAM) tool (v1.3), using default parameters and including the UniRef database option. The DNA GC content (% mol) of each isolate was calculated using the SeqinR package (96). Resistance Gene Identifier (RGI 5.2.0, CARD 3.1.2) was used to predict antimicrobial resistance genes within the 499 and 831b genomes (97). The mobility of the 831b plasmid was predicted using MOB-Typer (v3.1.0) (98, 99). Butyryl-CoA:acetate CoA-transferase and butyrate kinase genes were annotated with Bakta using the light database (v1.7) (100). The Butyryl-CoA:acetate CoA-transferase amino acid sequences were aligned with MAFFT using the L-INS-i method (default parameters, version 7) (101) and a tree was constructed using IQ-TREE with the LG+G4 model chosen based on Bayesian information criterion (v2.2.0) (93, 94). The tree was visualized with iTOL (95).

### Substrate utilization profiles

Substrate utilization analysis was performed using a Bioscreen C Analyzer (Growth Curves USA) in an anaerobic chamber. A basal liquid medium with or without the singular carbohydrate to be tested (1% w/v) was prepared according to a formulation described by Trachsel *et al*. (87). Strains were inoculated in basal media alone and then supplemented with individual substrates in replicates of nine. Optical density measurements (600 nm) were taken every hour for 48 h. Statistically significant increases in terminal and/or peak OD^600^ readings in the presence of a given substrate were indicative of an improvement in growth compared to basal medium (paired t-test; *P* < 0.05 considered significant). API 20A diagnostic strips were inoculated and interpreted according to the manufacturer’s specifications (bioMérieux) as an additional biochemical characterization measure. Media contained in the kit was supplemented with sodium acetate at a final concentration of 33 mM, a necessary component for adequate growth of both isolates. Test strips were incubated for 24-36 h at 39°C.

### Cellular fatty acid composition

The cellular fatty acid methyl ester (FAME) compositions of the isolates were analyzed with GC-MS using Agilent Technologies Model 6890 Gas Chromatograph coupled to a Model 5973 Mass Selective Detector capable of electrical ionization (EI) after approximately 36 h of growth in basal medium supplemented with 1% glucose at 39°C.

### Analysis of fermentation products

HPLC analysis of the SCFA fermentation products of 499 and 831b from glucose was performed using Aminex HPX-87H Column #1250140 (Bio-Rad). Sample tubes contained one mL of supernatant from a 24 h culture. Three separate cultures of each isolate were used for replication. Samples were injected into the column at a flow rate of 0.6 mL/min of 0.1N H^2^SO^4^, and UV detection was set to 210 nm. All values were normalized against a set of standards. SCFA concentrations >1 mM were considered positive. The SCFA concentrations (mM) were averaged across replicates.

### Testing for spore formation

The potential for spore formation from the genomic sequences of isolates 499 and 831b was predicted with Traitar (v1.1.1) (78). The isolates were subsequently tested for sporulation ability *in vitro* using several selection assays. Bacterial cells were grown in RTY broth for 48, 96, and 168 h in an anerobic chamber. An aliquot of broth culture from each timepoint was treated with an equal volume of ethanol (40 and 70%) at room temperature under ambient aerobic conditions for four hours to kill vegetative cells (102). Ethanol-treated cells were pelleted, washed, resuspended in PBS, and serially diluted before plating on basal medium with glucose, and RTY agar with and without 0.1% porcine bile extract, which served as a potential nutrient germinator (79). We evaluated the oxygen sensitivity of 499 and 831b and the potential for oxygen exposure to induce spore formation by incubating pure cultures in RTY broth for 48 h and then plating a dilution series on RTY agar. The plates were incubated under aerobic conditions at room temperature for different durations (24, 48, 96 h) before being returned to the anaerobic chamber incubator set to 39°C for 7 days.

### Microscopy techniques

Cells were prepared for electron microscopy from a 36 h culture in RTY media. For scanning electron microscopy visualization, bacterial cell cultures were put through a 0.22 µm Swinney filter, and the filters were fixed in 2.5% glutaraldehyde in 0.1 M Cacodylate buffer. The samples were stained by sequential exposure to osmium and thiocarbohydrazide (103), dehydrated through graded alcohols, and chemically dried with hexamethyldisilizane (104). Samples were decorated with a thin coating of gold/palladium mixture and viewed on the Hitachi TM3030Plus SEM. A negative stain was also performed on the cells. A 10 µl aliquot of culture was taken directly from each media tube and placed directly on a copper 200 mesh formvar carbon film grid (Electron Microscopy Sciences). After 60 seconds, the sample was overlaid with 2.0% Phosphotungstic acid (PTA, pH 7.0), for three seconds. The cultures were viewed with a ThermoFisher FEI Tecnai G^2^ BioTWIN electron microscope (FEI Co.) and images were taken with a side mount ORCA-HR camera (Advanced Microscopy Techniques).

### Metagenomic sequencing and analysis

We conducted metagenomic sequencing of DNA isolated from swine fecal and cecal samples that were part of a previously published study (80). The primary focus of the prior study (referred to above as *Salmonella* challenge study) was to assess the effects of feed amendments on the shedding of *Salmonella* following inoculation with *Salmonella* I 4,[5],12:i:-(SX 240). Fecal and cecal samples collected from pigs fed the Control diet or 5% RPS amended diet (MSP Starch Products Inc.) collected at 0, 2, 7, 14, and 21 DPI were used for metagenomic sequencing and SCFA measurements. Fecal SCFAs were quantified as described previously (80). DNA from fecal and cecal samples were extracted with the Qiagen MagAttract PowerMicrobiome Kit. Metagenomic sequencing libraries were prepared using a published modification of the Illumina DNA Prep kit protocol known as Hackflex (105) with the IDT for Illumina UD Indexes. The resulting libraries were pooled and sequenced on an Illumina NovaSeq 6000 with an S2 flow cell to generate 100 bp paired-end reads. Fecal and cecal metagenomic sequencing reads were mapped to the complete 831b and 499 genomes with bowtie2 (v2.5.1) (106). RPKM and genome coverage metrics were calculated using CoverM (v0.6.1) (107). The 831b or 499 strains were considered “present” in each metagenome if the fraction of genome covered by at least one read was >0.5, as has been suggested previously (81, 82). Differences in abundance (RPKM) and genome coverage between treatment groups were evaluated using the Wilcoxon Rank Sum Test implemented in R.

## Data availability

The previous assemblies of 499 (GCA_001940225.1) and 831b (GCA_001940165.1) have been updated. Oxford Nanopore sequencing data for the isolates 499 and 831b has been deposited under BioSamples SAMN37352064 and SAMN37352065, respectively, in BioProject PRJNA341691. All raw metagenomic sequence data has been deposited in the SRA and is available under BioProject PRJNA638426.

## Acknowledgements

We gratefully acknowledge Torey Looft for providing helpful discussion and expertise, Laura Hansen for technical assistance, and Judith Stasko of Microcopy Services at the National Animal Disease Center for assistance with imaging. This research used resources provided by the SCINet project and the AI Center of Excellence of the USDA Agricultural Research Service, ARS project number 0500-00093-001-00-D. This research was supported by appropriated funds from USDA-ARS CRIS project 5030-3200-225-00D and 5030-32000-227 and an appointment to the Agricultural Research Service (ARS) Research Participation Program administered by the Oak Ridge Institute for Science and Education (ORISE) through an interagency agreement between the U.S. Department of Energy (DOE) and the U.S. Department of Agriculture (USDA). ORISE is managed by ORAU under DOE contract number DE-SC0014664. All opinions expressed in this paper are the authors’ and do not necessarily reflect the policies and views of USDA, ARS, DOE, or ORAU/ORISE. We declare that the research was conducted in the absence of any commercial or financial relationships that could be construed as a potential conflict of interest. Mention of trade names or commercial products in this article is solely for the purpose of providing specific information and does not imply recommendations or endorsement by the U.S. Department of Agriculture. The USDA is an equal opportunity provider and employer.

---

**Figure S1.**
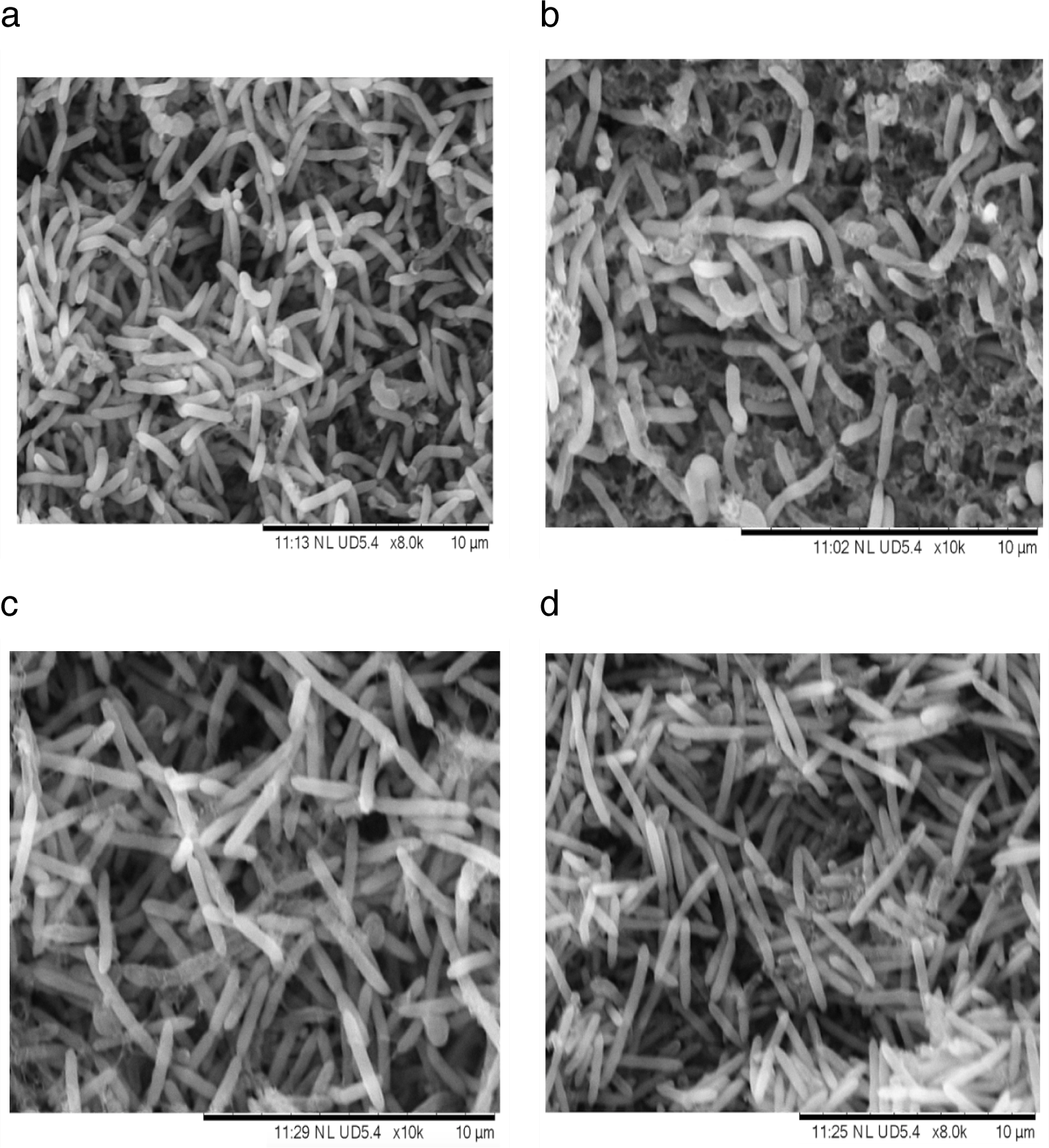
Scanning electron micrographs of isolate 831b (a,b) cells and 499 (c,d) cells. Cells were visualized in RTY medium following 36 h of incubation at 39°C.

**Figure S2.**
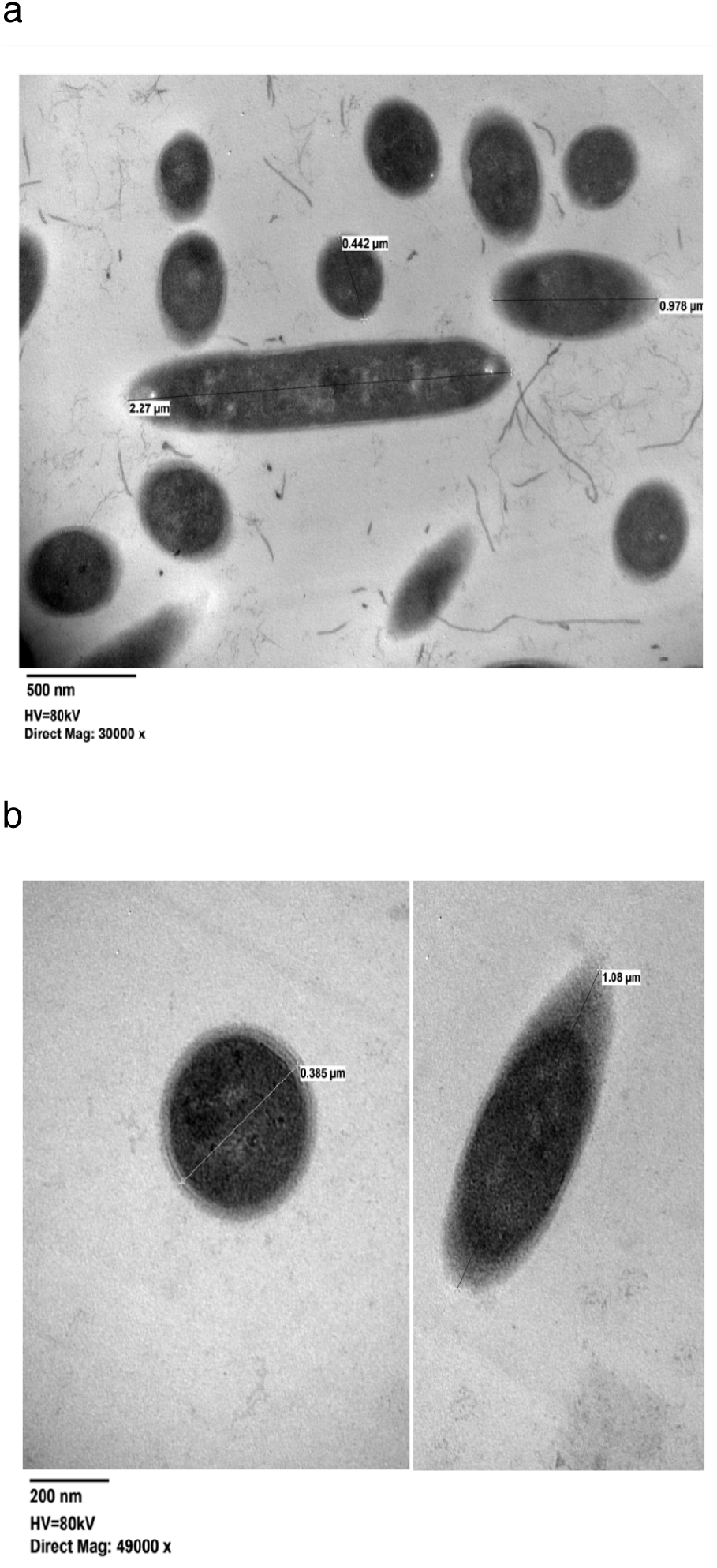
Transmission electron micrograph of isolate 831b (a) and 499 cells (b). Length and diameter of several cells are provided. Cells were visualized in RTY medium following 36 h of incubation at 39°C.

**Figure S3.**
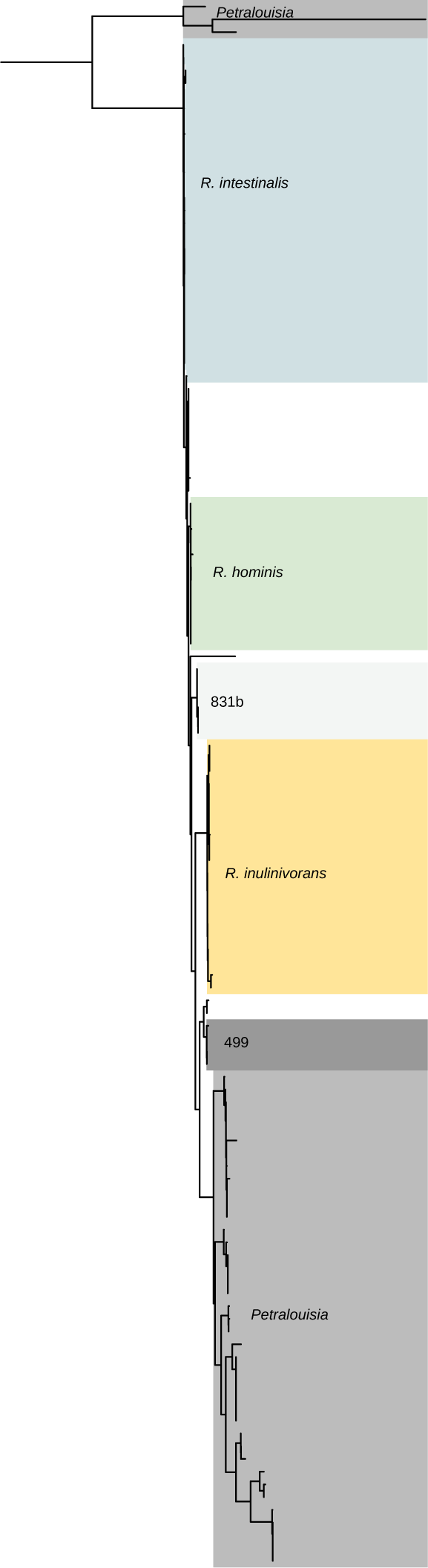
Maximum-likelihood tree based on the alignment of full-length 16S rRNA gene sequences of isolates 499 and 831b in addition to reference *Roseburia* and *Petralouisia* strains from GTDB. The tree is rooted at the midpoint and clades are labeled and highlighted with *Petralouisia* in gray, 831b in light gray, 499 in dark gray, *R. inulinivorans* in yellow, *R. hominis* in green, and *R. intestinalis* in blue.

**Figure S4.**
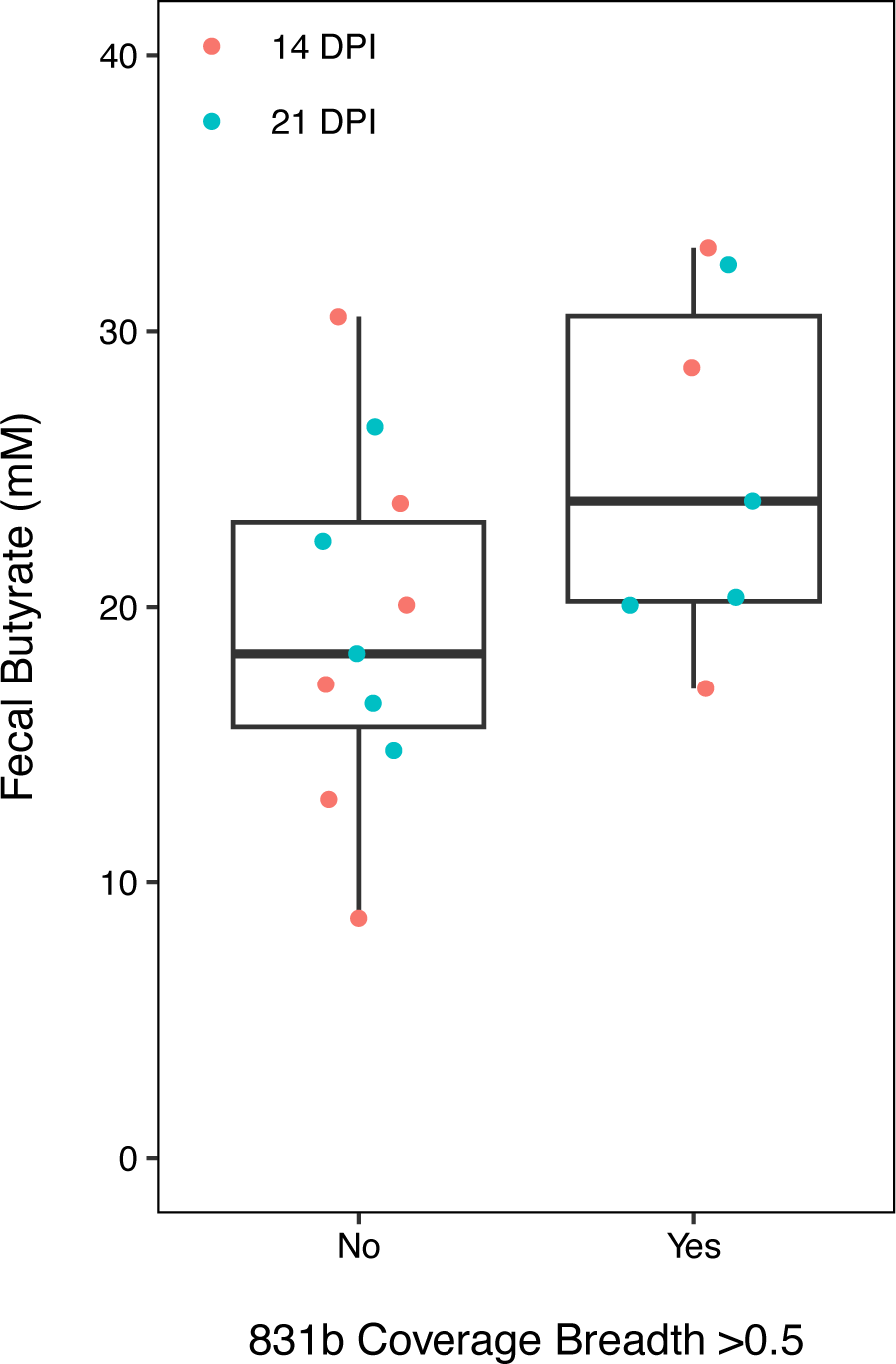
Fecal butyrate concentrations in samples from 14- and 21-days post inoculation in which 831b was determined to be present (coverage breadth >0.5) or not present (coverage breadth <0.5) in the metagenome. A coverage breadth of 0.5 indicates at least 50% of the genome had at least 1X coverage and has been suggested as a threshold for the assessing the presence of a strain in a metagenomic dataset (81, 82).

